# Swimming ability and flagellar motility of sperm packets of the volvocine green alga *Pleodorina starrii*

**DOI:** 10.1101/2023.06.08.544266

**Authors:** Azusa Kage, Kohei Takahashi, Hisayoshi Nozaki, Tetsuya Higashiyama, Shoji A. Baba, Takayuki Nishizaka

## Abstract

Eukaryotic flagella collectively form metachronal waves that facilitate the ability to cause flow or swim. Among such flagellated and planktonic swimmers, large volvocine genera such as *Eudorina*, *Pleodorina* and *Volvox* form bundles of small male gametes (sperm) called “sperm packets” for sexual reproduction. Although these sperm packets reportedly have flagella and the ability to swim, previous studies on volvocine motility have focused on asexual forms and the swimming characteristics of sperm packets remain unknown. However, it is important to quantify the motility of sperm packets and sperm in order to gain insights into the significance of motility in the sexual reproduction of planktonic algae. In this study, we quantitatively described the behavior of three flagellated forms of a male strain of *Pleodorina starrii*—asexual colonies, sperm packets, and single dissociated sperm—with emphasis on comparison of the two multicellular forms. Despite being smaller, sperm packets swam approximately 1.4 times faster than the asexual colonies of the same male strain. Body length was approximately 0.5 times smaller in the sperm packets than in asexual colonies. The flagella from sperm packets and asexual colonies showed asymmetric waveforms, whereas those from dissociated single sperm showed symmetric waveforms, suggesting the presence of a switching mechanism between sperm packets and dissociated sperm. Flagella from sperm packets were approximately 0.5 times shorter and had a beat period approximately twice as long as those from asexual colonies. The flagella of sperm packets were densely distributed over the anterior part of the body, whereas the flagella of asexual colonies were sparse and evenly distributed. The distribution of flagella, but not the number of flagella, appear to illustrate a significant difference in the speeds of sperm packets and asexual colonies. Our findings reveal novel aspects of the regulation of eukaryotic flagella and shed light on the role of flagellar motility in sexual reproduction of planktonic algae.

## Introduction

Collective motion is common in nature, such as that observed in schools of fish [1], flocks of birds [2] or herds of sheep [3]. Movements in fluids such as swimming of fish and flying of birds can be characterized by the Reynolds number *Re*, which is defined as *Re* = ρ*vL*⁄μ, where *ρ* is the density of fluid, *v* is the speed, *L* is the characteristic length (e.g., body length), and *µ* is the viscosity of the fluid. The Reynolds number represents how inertia and viscosity dominate the movement; higher *Re* indicates the movement is more affected by inertia than by viscosity and vice versa. For low *Re* (<<1), where the spatial scale is small (< 1 mm), inertia does not contribute, and viscosity dominates the movement. Striking examples of low-*Re* movement are microbial swimming and flagellar/ciliary motion [4,5]. Metachronal waves of eukaryotic flagella or cilia [6,7] are a form of collective motion of many flagella/cilia, and are important for microbial swimming and flow generation inside the animal body. Understanding such collective motion on a small scale would lead to better understanding of the ecology of planktonic organisms that swim with flagella/cilia and manipulation of small-scale movements such as those in drug delivery systems.

Volvocine green algae, ranging from unicellular *Chlamydomonas* [8] and four-celled *Tetrabaena* [9] to thousands-celled *Volvox* [10,11], are planktonic swimmers with flagella that have been used as model organisms to investigate the dynamics of multiple flagella. They have two equal-length flagella in one cell, and flagella in the entire colony have been shown to move collectively in many cases. Previous studies on the motility of multicellular volvocines focused on asexual colonies that are in their usual form in culture. Although they have been known to sexually reproduce [12] and the sex-determining locus have been identified in several species (*Pleodorina starrii* [13], *Volvox carteri* [14], *Gonium pectorale* [15], *Yamagishiella unicocca* and *Eudorina* sp. [16]), the function of motility in sexual reproduction in multicellular volvocines remains unclear.

In the process of sexual reproduction, bundles of small male gametes or sperm called “sperm packets” have been reported in large multicellular volvocine genera such as *Eudorina* [17], *Pleodorina* [18,19], and *Volvox* [20,21]. Male gametes or sperm in these genera are characterized as biflagellated cells with elongated cell bodies, which differs from typical animal sperm with a single flagellum [22,23]. Sperm are bundled as packets in male colonies, and these sperm packets swim into female colonies [18,19]. Sperm packets subsequently dissociate into single sperm after arrival at the female colonies and then fertilization occurs between dissociated single sperm and female gametes held by female colonies. However, these processes have only been described qualitatively, and no study has quantitatively characterized the movement of flagella in sperm packets or sperm in volvocines.

In this study, we focused on the motility of the three flagellated forms of a male strain of the volvocine alga *Pleodorina starrii* (Figure 1)—asexual colonies, sperm packets, and single dissociated sperm—with emphasis on the comparison of the two multicellular forms (asexual colonies and sperm packets). *P. starrii* has three sexes: male, female, and hermaphrodite [24]. We found that the swimming speed of sperm packets was approximately 1.4 times faster than that of asexual colonies, whereas the body size in sperm packets was approximately 0.5 times smaller than that in asexual colonies. A comparison with a previous computational study [25] suggested that the distribution of flagella over the body illustrates the difference in swimming speeds between sperm packets and asexual colonies. Furthermore, we found that flagella from single dissociated sperm showed symmetric motion, whereas flagella from sperm packets showed asymmetric motion, suggesting a waveform switching mechanism in sperm cells. Our findings provide insights into the function of motility in the sexual reproduction of volvocines, which had previously been overlooked.

**Fig. 1.**
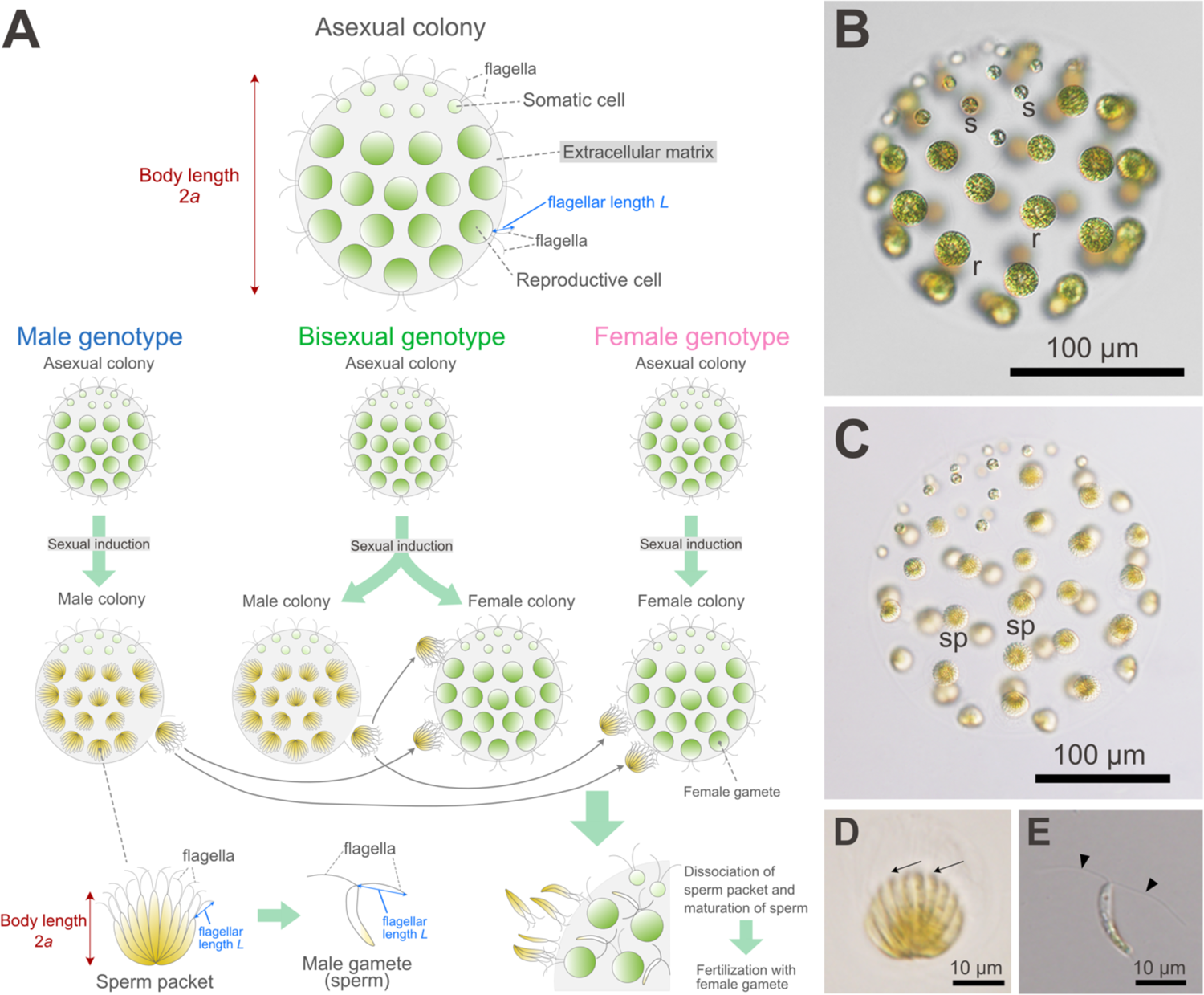
Light microscopic features of *Pleodorina starrii*, based on Takahashi et al. [24]. (A) Schematic representation of the life cycle. Colonies proliferate asexually. Each colony consists of 32–64 cells with equal-length flagella. The cycle of sexual reproduction begins under a lack of nitrogen, and consequently, male colonies with sperm packets—bundles of male gametes (sperm) with two equal-length flagella—are formed in male and bisexual phonotypes. The sperm packet swims to a female colony produced in female or bisexual phenotypes, and dissociates into sperm that fertilize with female gametes within the female colony. The body length of asexual colonies and sperm packets (2*a*) was measured as described in the Materials and Methods section. The flagellar lengths of asexual colonies, sperm packets, and dissociated single sperm (*L*) were measured as the length between the tip and base of the flagellum (see Materials and Methods). For details of the three sex phenotypes (male, female, and bisexual), see Takahashi et al. [24]. (B) 64-celled mature asexual colony. (C) Sexually induced male colony with sperm packets (sp). (D) Sperm packet. (E) Dissociated single sperm. All photographs and illustrations are original. For materials and methods of light micrographs and illustrations, see Takahashi et al. [24].

## Materials and Methods

### Culture

*Pleodorina starrii* was maintained at 20 °C in the artificial freshwater-6 (AF6) medium (pH 6.6) [26] under a 10:14 light/dark (LD) cycle. Asexual colonies were transferred into the Volvox thiamine acetate (VTAC) medium (pH 7.5) [26] and incubated for 2–3 days at 25 °C under a 12:12 LD cycle. Sperm packets were induced by transferring 3-day colonies from the VTAC medium into a mating medium that lacked a nitrogen source [18,24]. The unisex male strain P7 [24] (NIES-4480 [27]) was used unless otherwise stated. The dissociation process of the sperm packet was observed using the unisex male strain P7 × P10 F1–14, which is the offspring of P7. The experiments were performed at a temperature of approximately 23–25 °C during the subjective daytime.

### Recording of free swimming

Swimming activity was recorded using an upright microscope BX53 (Olympus, Tokyo, Japan) with a halogen lamp (Olympus, Tokyo, Japan), dry dark-field condenser (U-DCD, Olympus, Tokyo, Japan), and 620-nm long-pass filter to prevent phototaxis (SCF-50S-62R, OptSigma, Tokyo, Japan). Although phototaxis was not examined in the present study, the wavelength was chosen based on studies on *Chlamydomonas* phototaxis, in which it did not induce phototaxis [28]. *P. starrii* was confined between a glass slide (S1126, Matsunami, Osaka, Japan) and coverslip (18 × 18 mm, Matsunami, Osaka, Japan) with an ∼10 × 10 mm square hole in a 0.5 mm silicone rubber spacer (Kogugo, Tokyo, Japan). Movies were captured at 14.3 fps using a CMOS camera (UI-3250-CP, IDS, Obersulm, Germany) to monitor swimming and at 1,000 fps using a high-speed CMOS camera (VCC-H1540, Digimo, Tokyo, Japan) to monitor flagellar motility. A 10× objective lens (UPlanXApo, NA 0.40, Olympus, Tokyo, Japan) was used to observe sperm packets, whereas a 4× objective lens (UPlanFl, NA 0.13, Olympus, Tokyo, Japan) was used to observe asexual colonies. Swimming trajectories and body lengths were quantified using the TrackMate plugin [29] in ImageJ (Fiji). The swimming speed of each individual was calculated as the mean swimming speed along the trajectory. The TrackMate plugin estimates the diameter of objects (bright spots); here, the minimum value of the estimated diameter in the track was considered as the body length or twice the radius *a* [25] because the estimated diameter was the minimum when the object was in focus.

Flagellar motility during swimming was visualized using the same setup (objective 10×) except that a mercury lamp (Olympus, Tokyo, Japan) without a long-pass filter (i.e., white light) was used in order to ensure sufficient light to visualize the movement of flagella, of which diameter is ∼250 nm [30] and beating frequency is tens of Hz [31], which require high shutter speed (i.e., short exposure time).

### Recording of flagellar motility of sperm packets

To visualize flagellar movement, sperm packets were captured using a glass micropipette [32]. Glass pipettes with a tip diameter of ∼20 µm were fabricated with a puller (PC-100, Narishige, Tokyo, Japan) and microforge (MF2, Narishige, Tokyo, Japan) from the capillary (G-1, Narishige, Tokyo, Japan). The glass pipette was set to the injector (IM-11-2, Narishige, Tokyo, Japan) on an inverted microscope (IX71, Olympus, Tokyo, Japan); 0.5–2 µL droplets of the culture medium containing sperm packets were placed on a glass-bottom dish (P50G-0-30-F, Mattek, Ashland, MA, US), and the object inside the droplet was suctioned with the pipette. The droplets were covered with mineral oil (8042-47-5; ICN Biomedicals, Aurora, OH, US) to prevent evaporation during observation. Movies were captured at 1,000 fps using a high-speed CMOS camera (VCC-H1540, Digimo, Tokyo, Japan), long-working distance phase-contrast optics (Olympus, Tokyo, Japan) and a 40× objective lens (UPlanFl, NA 0.75, Olympus, Tokyo, Japan) with a halogen lamp (TH4-100, Olympus, Tokyo, Japan) and a 620-nm long-pass filter (SCF-50S-62R, OptSigma, Tokyo, Japan).

### Analysis of flagellar motility

The flagellar waveform was analyzed using the lab-made software Bohboh [33]. Semi-auto-tracking of flagella was applied to the dark-field images of asexual colonies and dissociated single sperm. For pipette-captured sperm packets, phase-contrast images of the flagella were tracked manually because of poor contrast. From the time series of flagellar traces, we obtained space–time plots (kymographs) of curvature, along with flagellar length and time [34]. Flagellar beat periods were visually estimated from the space–time plots, and flagellar lengths were calculated from traces on Bohboh. As the length fluctuated along the beat period owing to the three-dimensionality of the waveform, the mean length for each flagellum (50–100 frames) was adopted as the flagellar length.

### Immunofluorescence microscopy

To visualize the distribution of flagella, indirect immunofluorescence microscopy was performed according to Matsuura et al. [35], using the same method as that used for staining *Chlamydomonas* flagella. The samples were fixed in methanol. Antiacetylated α-tubulin (mouse monoclonal 6-11B-1, ab24610, Abcam, Cambridge, UK) diluted in blocking buffer (containing 10 mM phosphate buffer pH 7.2, 5% normal goat serum, 5% glycerol, 1% cold fish gelatin, 0.004% Na azide) to 1:500 was used as the primary antibody, and samples were incubated with the primary antibody at 37 °C for 1 hour. DyLight® 488-conjugated goat anti-mouse IgG (ab96871, Abcam, Cambridge, UK) diluted in blocking buffer to 1:500 was used as the secondary antibody, and samples were incubated with the secondary antibody at 37 °C for 1 hour. Nuclei were stained with 50 µg/mL Hoechst 33342 (Nacalai Tesque, Kyoto, Japan) in PBS for 10 minutes before immunofluorescence staining. Confocal images were captured using an FV-3000 microscope (Olympus, Tokyo, Japan) equipped with a 100× oil-immersion objective lens (UPlanApo, NA 1.50, Olympus, Tokyo, Japan) at the Department of Life Sciences, Gakushuin University. To grasp the three-dimensional structure of sperm packets and asexual colonies, Z-stacks were overlaid using the FV-3000 software (Olympus, Tokyo, Japan).

## Results

### Sperm packets swam faster than asexual colonies

To characterize the swimming of sperm packets, we recorded and compared the swimming of sperm packets with asexual colonies of the same male strain (P7) at low magnification. Figure 2A shows the relationship between swimming speed and body length of sperm packets (n = 185) and asexual colonies (n = 313). Hereafter, “body length of a sperm packet” indicates the size of the whole sperm packet, not a single cell (sperm). Sperm packets had smaller body lengths and higher swimming speeds than asexual colonies. The top and right panels of Figure 2A show the distribution of the body lengths and swimming speeds of sperm packets and asexual colonies. The body lengths of sperm packets and asexual colonies were 24.7 ± 5.9 µm (mean ± standard deviation [S.D.], n = 185) and 54.7 ± 14.9 µm (mean ± S.D., n = 313), respectively. The swimming speeds of sperm packets and asexual colonies were 308 ± 73 µm/s (mean ± S.D., n = 185) and 215 ± 63 µm/s (mean ± S.D., n = 313), respectively, indicating that sperm packets swam approximately 1.4 times faster than asexual colonies. Both the speed and the body length were significantly different between sperm packets and asexual colonies (speed: *p* < 10^-36^ and body length: *p* < 10^-74^ in Mann–Whitney *U*-test). The speed and body length showed no correlation in sperm packets (*R* = 0.07, *p* = 0.34 in Pearson’s correlation test) and a weak but significant negative correlation in asexual colonies (*R* = −0.31, *p* < 10^-7^). Figures 2B and 2C show the swimming trajectories of the sperm packets and asexual colonies, respectively. Both exhibited nearly straight trajectories within an observation field of less than 1 mm.

**Fig. 2.**
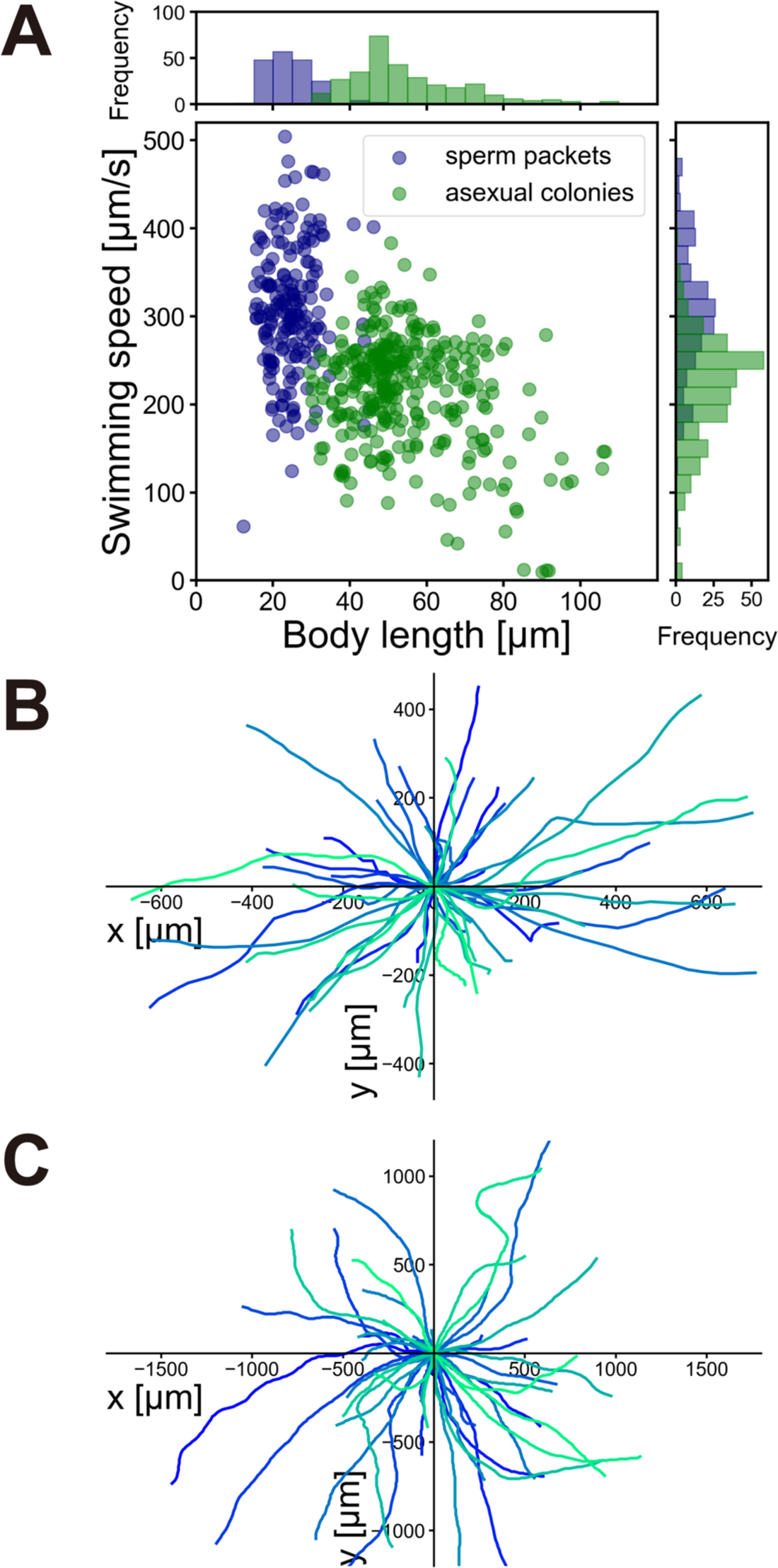
Swimming of sperm packets and asexual colonies. (A) Swimming speed plotted against body length (diameter). The top panel shows a histogram of body length; the right panel shows a histogram of swimming speed (navy: sperm packets, green: asexual colonies). (B) Swimming trajectories of sperm packets for 50 randomly chosen trajectories. The duration of tracking was 1.30 ± 0.69 sec (mean ± S.D.). (C) Swimming trajectories of asexual colonies for 50 randomly chosen trajectories. The duration of tracking was 4.21 ± 3.33 sec (mean ± S.D.).

### Visualization of flagella in dark-field microscopy

To clarify the reason for the difference in speed between sperm packets and asexual colonies, we visualized their flagellar movements. Figure 3 and Supplementary Movie S1 show a swimming sperm packet viewed from the side using a dark-field microscope. The sperm packet swam with flagella anterior (i.e., sperm packets are puller-type swimmers [36]) and metachronal waves of the flagella traveling from the top region to the periphery of the packet, although it was difficult to determine the relationship between the direction of the wave and the power stroke of the flagella because of the dense distribution of flagella. The flagella of asexual colonies (Figure 4, Supplementary Movie S2) were evenly distributed over the body, and the metachronal waves were symplectic (i.e., waves propagated in the same direction as the power stroke of the flagella [37]), as observed in *Volvox* [38].

**Fig. 3.**
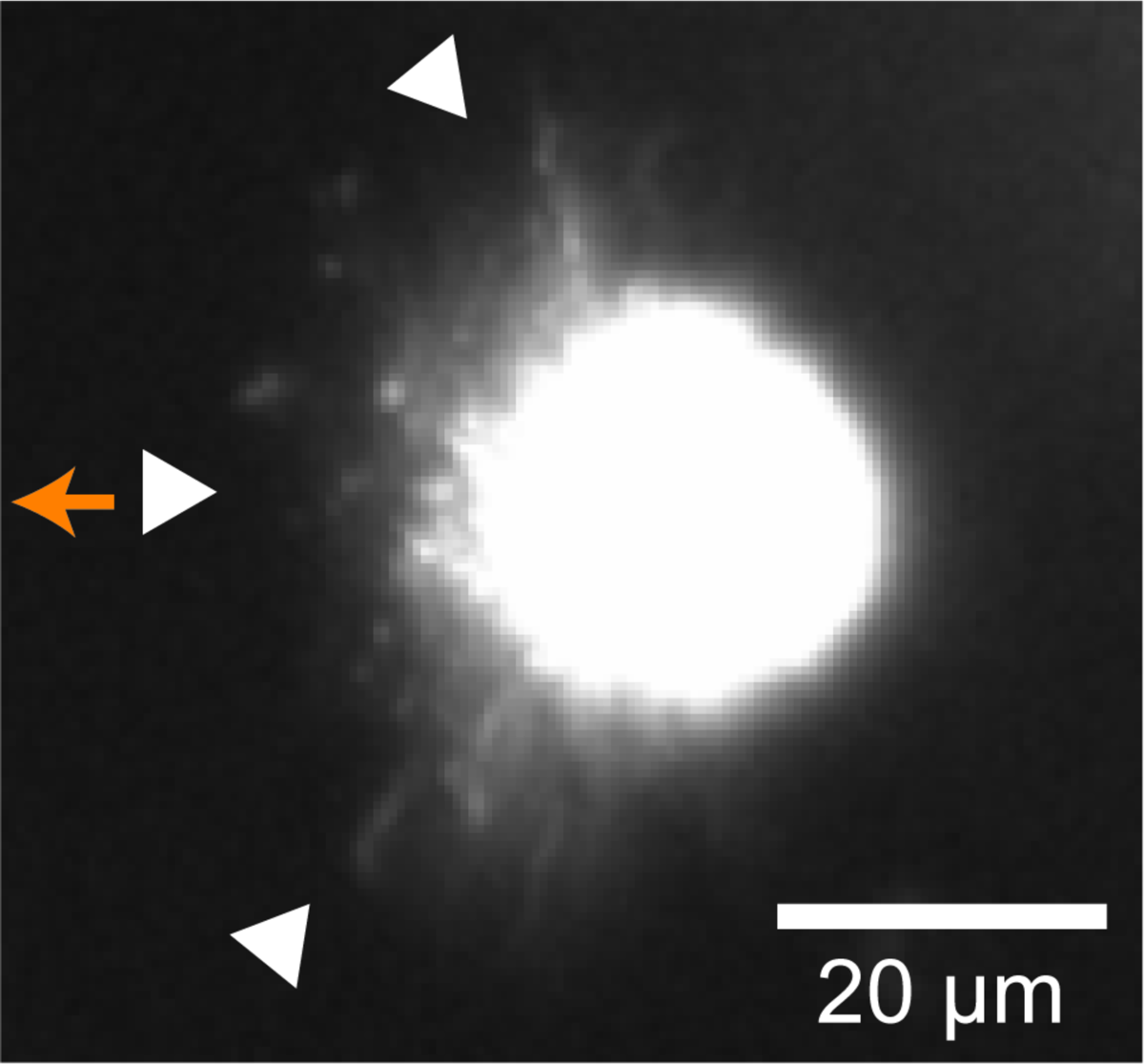
Image of a sperm packet (seen from the side) under dark-field microscopy. The sperm packet moved using anterior flagella (i.e., puller-type swimming). The metachronal wave propagated from the center (see Supplementary Movie 1). White triangles indicate flagella and the orange arrow indicates the swimming direction.

**Fig. 4.**
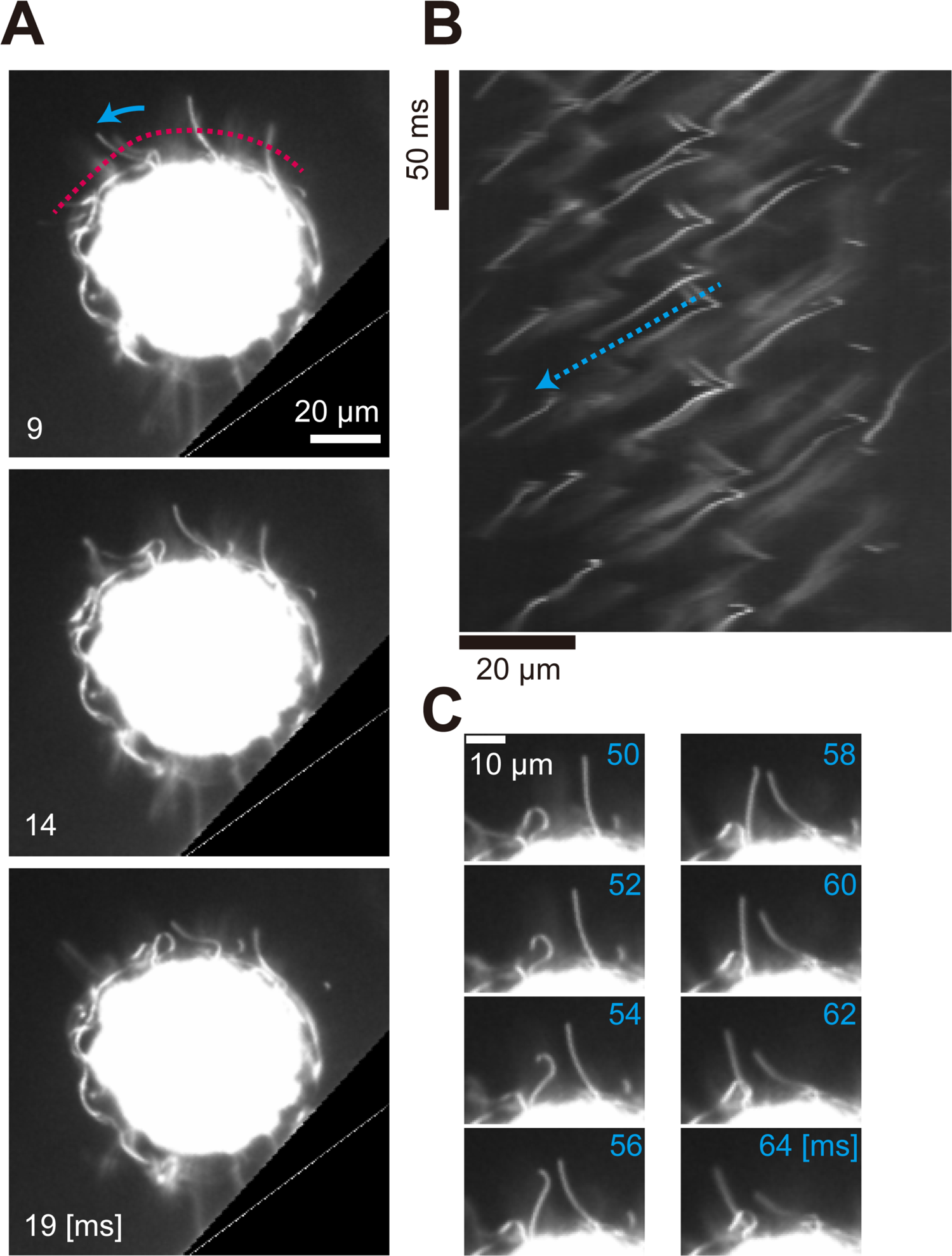
An asexual colony in dark-field microscopy. (A) Images of an asexual colony. Flagella beat with asymmetric waveform. Registered images of a rolling colony near the wall are shown every 5 ms. The image of the silicone rubber spacer (wall) was deleted (black triangle in the lower right) for proper registration (see Supplementary Methods). The blue arrow indicates the direction of the effective stroke; the broken red line indicates where the kymograph (B) was drawn. Time [ms] corresponds to the time axis in (B). (B) Kymograph of the movement of flagella drawn from the broken red line in (A). The broken blue arrow indicates the propagation of metachronal waves. (C) Flagella showing symplectic metachronal waves. Both the effective stroke of the flagella and the metachronal wave propagated from the left to the right side of the image.

Sperm packets disassemble into individual sperm after arrival in female colonies [19]. Figure 5 and Supplementary Movie S3 show the process of disassembling a sperm packet, as recorded by dark-field microscopy. The packet disassembled gradually as the sperm moved randomly. To date, the trigger for this dissociation has not been identified. Figure 6 and Supplementary Movie S4 show a single sperm after disassembly of a packet; the two flagella of the sperm beat alternately, and individual sperm barely propel themselves. Flagellar movement was symmetric (i.e., flagella bent in two directions [39]), similar to human or sea urchin sperm.

**Fig. 5.**
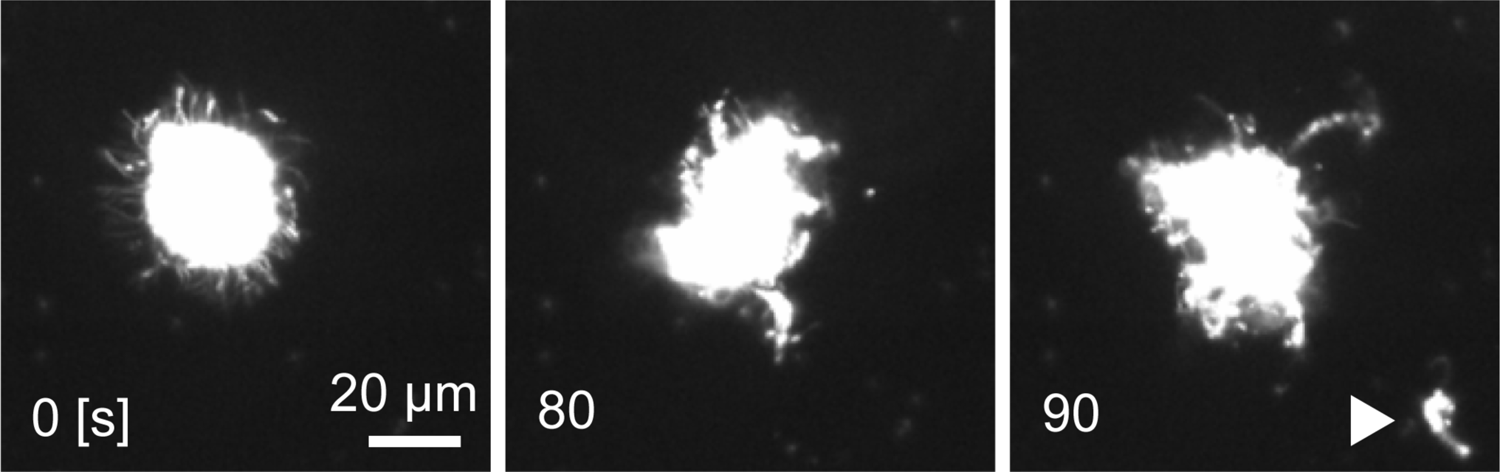
Dissociation of a sperm packet viewed using dark-field microscopy. The sperm packet gradually collapsed, and the dissociated sperm jumped out of the packet. The white triangle in 90 sec indicates dissociated sperm.

**Fig. 6.**
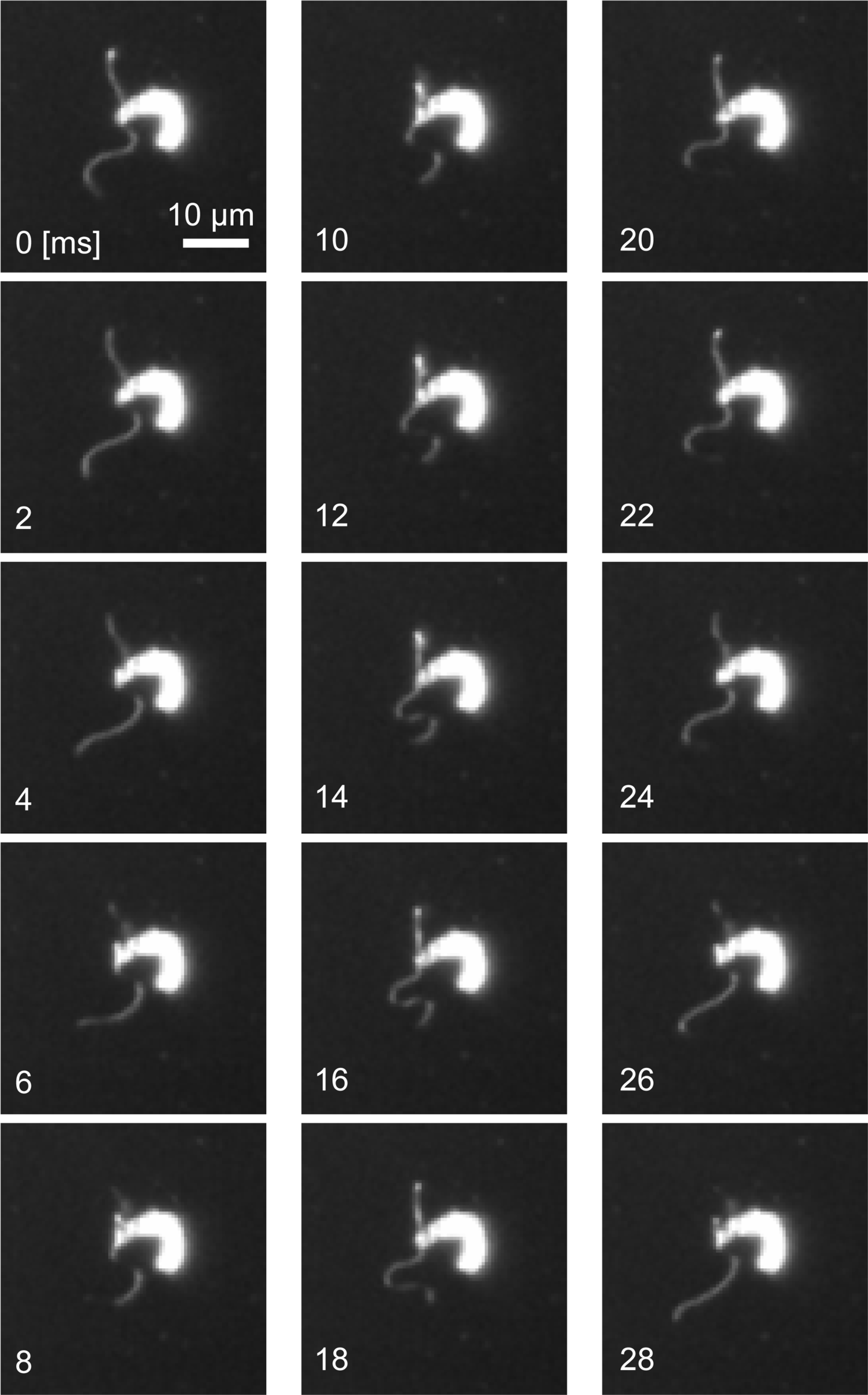
Dark-field microscopy images of a single dissociated sperm. The two flagella beat with a symmetric waveform and did not synchronize. One beat period was approximately 25 ms. Images are shown every 2 ms.

### Visualization of flagella of sperm packets held on the micropipette

As it was difficult to analyze flagellar motility from freely moving sperm packets (Figure 3), we suctioned the sperm packets into a glass micropipette. Figure 7 and Supplementary Movie S5 show the flagellar motility of a sperm packet held on a micropipette, using phase-contrast microscopy. We found that the flagella beat with an asymmetric waveform (i.e., flagella bent in only one direction). The effective stroke was directed from the top (anterior) region of the body to the periphery, and the metachronal waves were also directed from the top to the periphery (Figure 7B, C). Thus, the metachronal waves in the sperm packets were symplectic, similar to those in asexual colonies.

**Fig. 7.**
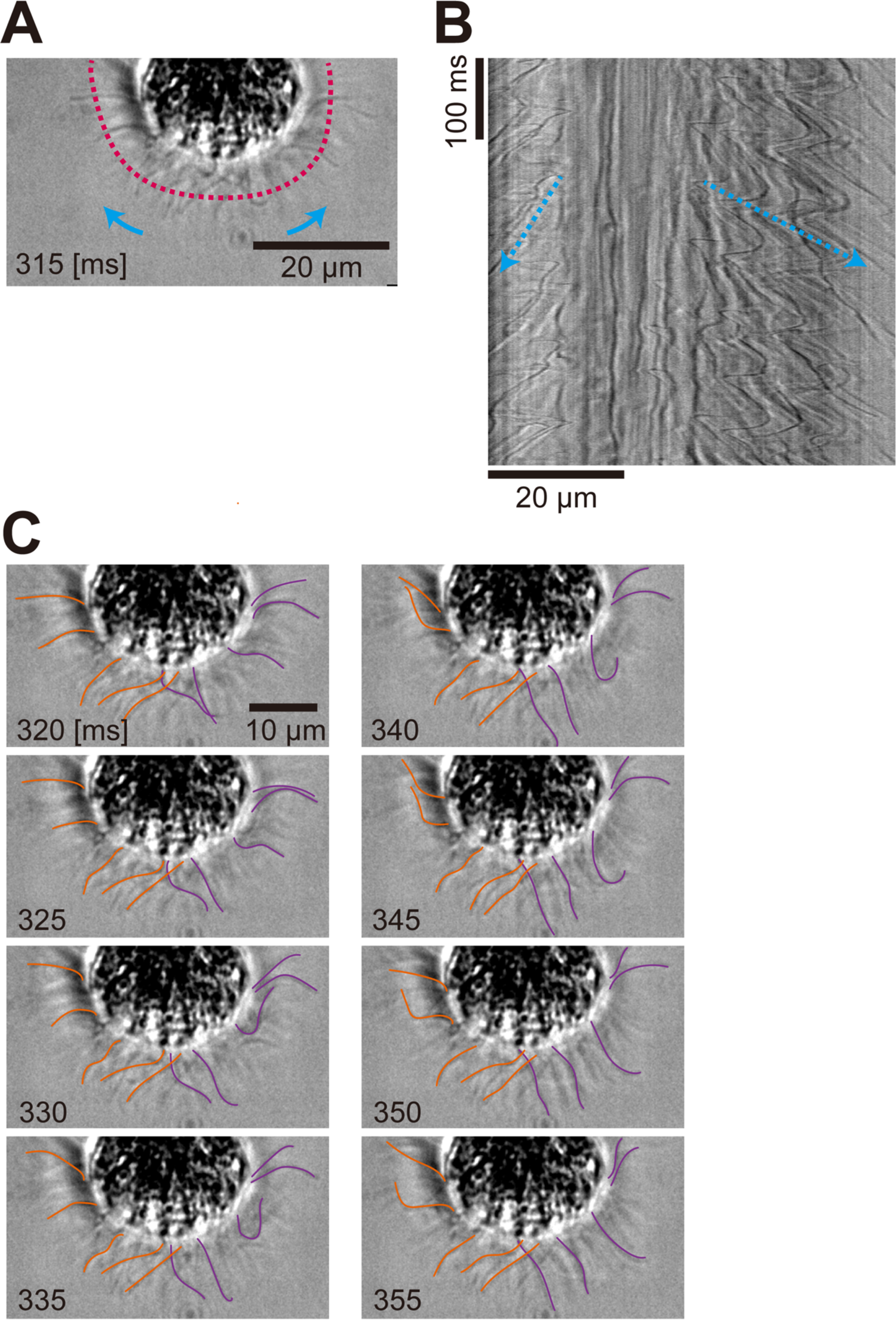
Sperm packet captured by a micropipette in phase-contrast microscopy. (A) Image of the sperm packet. The flagella beat with an asymmetric waveform. Images are shown every 5 ms. Blue arrows indicate the direction of effective stroke, and the broken red line indicates where the kymograph (B) was drawn. Time [ms] corresponds to the time axis in (B). Contrast enhanced. (B) Kymograph of the movement of flagella drawn from the red broken line in (A). Blue broken arrows indicate the propagation of metachronal waves. (C) Traces of flagella showing metachronal waves (purple: waves to the right side, orange: waves to the left side). Only a fraction of flagella were traced manually. Time [ms] corresponds to the time axis in (B).

### Analysis of flagellar waveforms

We tracked and quantified flagellar movement. Figures 8A, B, and C show a time series of flagellar waveforms traced from an asexual colony (Figure 4), dissociated single sperm (Figure 6), and sperm packets (Figure 7), respectively. The flagellum from the asexual colonies (Figure 8A) showed asymmetric waveforms, bending in one direction, similar to the colonies of other Volvocales [11]. The flagellum from the dissociated single sperm (Figure 8B) showed symmetric waveforms with principal and reverse bends [39]. The flagellum from sperm packets (Figure 8C) showed asymmetric waveforms with a usually larger period. When we calculated the mean curvature over a beat period at the midpoint of the flagella, the asexual colony (Figure 8A) showed a curvature of 0.11 rad/µm (at 10 µm from the base), the dissociated single sperm (Figure 8B) showed 0.03 rad/µm (at 7 µm from the base), and the sperm packet (Figure 8C) showed 0.08 rad/µm (at 5 µm from the base). The flagellum appeared shorter in sperm packets than in single sperm or colonies. As shown in Figure 8D–F, space–time plots of curvature along with time and flagellar length supported these views; Figures 8D and 8F indicate that the flagella from the asexual colony and the sperm packet bent in only one direction (i.e., asymmetric waveform), whereas Figure 8E shows that the flagella from the single dissociated sperm bent in two directions (i.e., symmetric waveform).

**Fig. 8.**
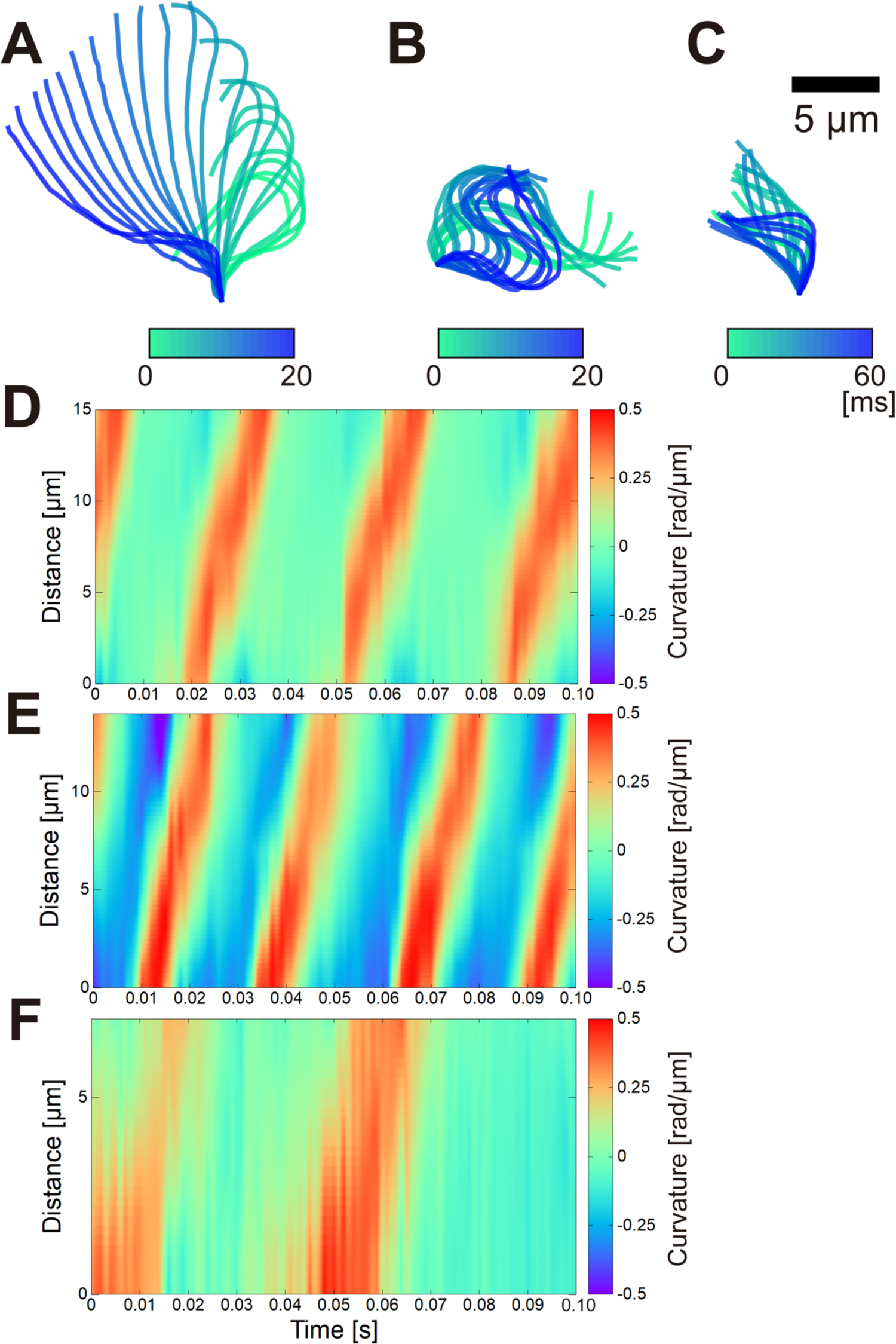
Flagellar waveform of (A) an asexual colony, (B) single dissociated sperm, and (C) a sperm packet. (A) and (B) show 20 frames at 1,000 fps, and (C) shows 60 frames every 3 frames. (D)–(F) Space–time plots of curvature along with time and flagellar length obtained from (A)–(C). Asymmetric waveforms in (D) and (F) show stripes with curvatures of positive (red) and near-zero (green) values, indicating a one-directional bend, whereas the symmetric waveform in (E) shows stripes of positive (red) and negative (blue) curvature, indicating a two-directional bend.

Using waveform analysis (Figure 8, Supplementary Figure S1 and S2), we estimated the beat period and length of the flagella (Supplementary Table 1). The beat period was 36 ± 13 ms in asexual colonies (mean ± S.D., n = 4), 30 ± 10 ms in single dissociated sperm (n = 3), and 69 ± 18 ms in sperm packets (n = 3). The flagellar length was 22.1 ± 7.8 µm in asexual colonies (n = 4), 16.7 ± 1.2 µm in single dissociated sperm (n = 3), and 11.0 ± 0.5 µm in sperm packets (n = 3).

### Visualization of the distribution of flagella

Indirect immunofluorescence microscopy was used to clearly visualize the distribution of flagella. Figures 9A and 9B show the distribution of flagella (green) in sperm packets and asexual colonies, respectively. The flagella of sperm packets were densely distributed over the anterior part of the body, whereas the flagella of asexual colonies were sparse and evenly distributed.

**Fig. 9.**
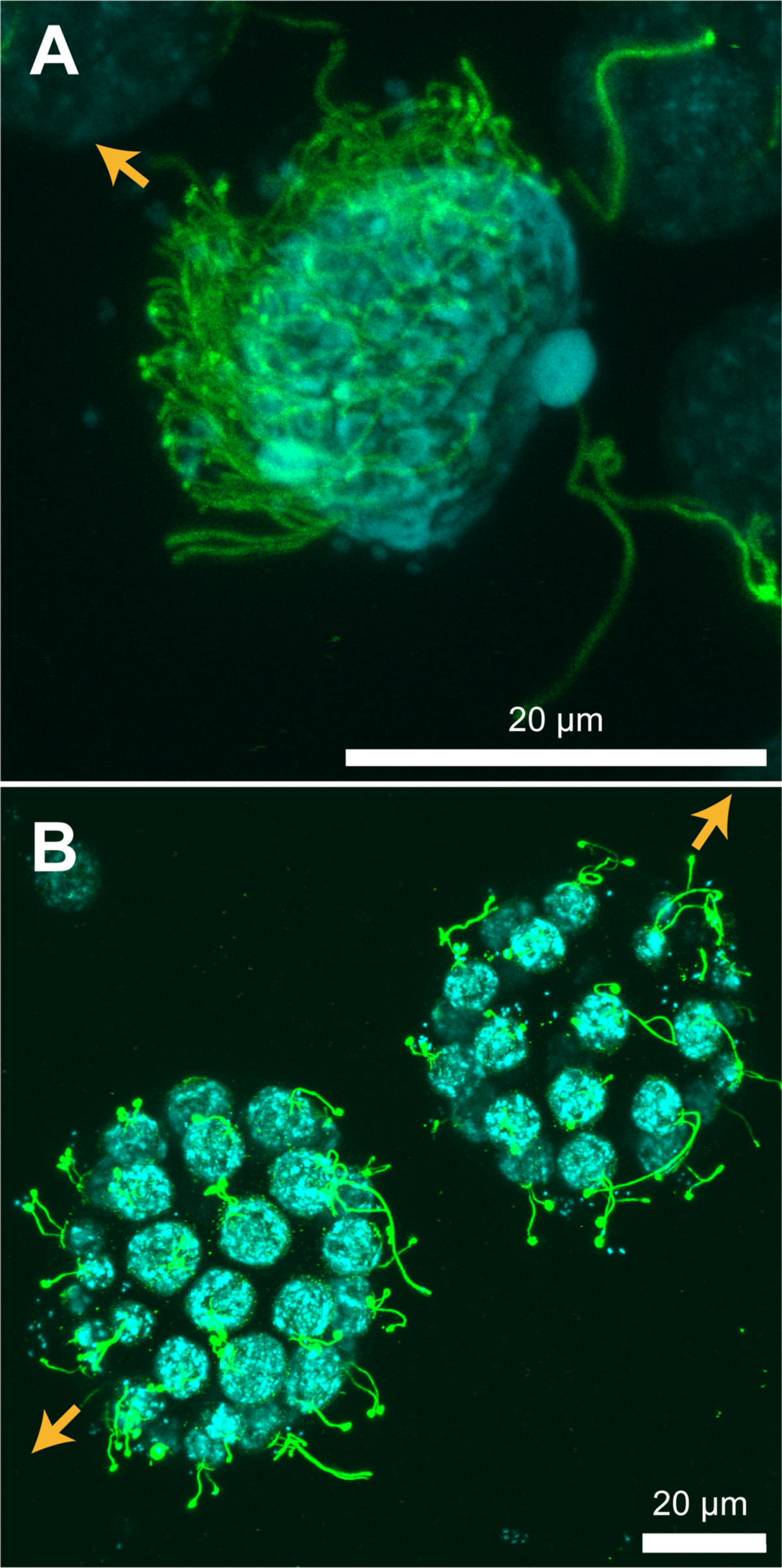
Distribution of flagella in immunofluorescence microscopy. (A) Sperm packet; (B) asexual colonies. Green structures represent acetylated α-tubulin (flagella) and blue structures represent nuclei. Z projections of confocal images are shown. Orange arrows indicate the swimming direction. Note that flagella were densely distributed in the sperm packet, whereas the asexual colony had sparse distribution of flagella.

## Discussion

### Mechanisms of straight swimming in asexual colonies and sperm packets

Both asexual colonies and sperm packets showed nearly straight swimming in a field of view of less than 1 mm (Figure 2B, C). Although both showed straight trajectories, the mechanisms are thought to be different, because the distribution (Figure 9) and metachronism (Figure 4, Figure 7) of flagella differ between asexual colonies and sperm packets. In asexual colonies, the distribution of flagella is nearly uniform, and each pair of flagella is nearly identical over a spherical body, although there might be anterior–posterior differences as observed in *Volvox* [11]. Therefore, the time-averaged force distribution generated by flagella is supposedly uniform over the asexual colony (i.e., neutral swimming), and this results in straight swimming. In other neutral swimmers [40] such as the ciliates *Paramecium* [41] and *Tetrahymena* [42], helical trajectories are frequently observed possibly because the metachronal waves are oblique against the body axis [41,43] or the cilia around the mouth (oral groove) move differently from the cilia in the other parts [41]; however, these were not the case in the asexual colonies of *P. starrii* in the present study. In contrast, sperm packets were pullers with tens of flagella in the anterior part (Supplementary Movie S1). The forces generated by flagella are stronger in the anterior part even if averaged over time. Symplectic metachronal waves are generated from the anterior to the posterior part of the packet (Figure 7, Supplementary Movie S5), resulting in straight swimming in the a–p axis direction.

### The normalized body length is similar, but the normalized speed is dramatically different between sperm packets and asexual colonies

Table 1 summarizes the swimming parameters of the sperm packets and asexual colonies. We found that, on average, sperm packets had approximately 0.5 times smaller body length, 1.4 times higher speed, 0.5 times shorter flagellar length, and twice longer beat period than sperm packets. Asexual colonies contained 32–64 cells [19] whereas the sperm packets contained 20–50 cells (n = 3). Thus, it is reasonable to assume that the number of flagella is comparable between the asexual colonies and sperm packets. From the numerical simulation of ciliated spherical bodies, Omori et al. [25] showed that the normalized swimming speed Ū/(L/T), where Ū is the swimming speed, *L* is flagellar length, and *T* is flagellar beat period, decreased by a power of −2 of the normalized body length 2*a*/*L* (where *a* is the radius of the body) if the number of the flagella *N* was fixed for a normalized body length ranging from 10 to 30. If we consider the mean flagellar length of each category, sperm packets and asexual colonies had normalized body lengths of 2.2 and 2.5, respectively (i.e., quite similar). However, the normalized swimming speeds were dramatically different between the two categories. If the normalized body size and direction of the metachronal waves (symplectic or antiplectic) were the same, the number of flagella would cause a difference in swimming speed (i.e., a smaller number of flagella results in a slower speed) [25]. However, the number of flagella would not differ in our case. Thus, in our study, the difference in speed could be attributed to the difference in the distribution of flagella (Figure 9), which was not considered by Omori et al. [25]. As the cell shape is elongated in sperm packets but spherical in asexual colonies [18,19,24], the spacing of the flagella would be smaller in sperm packets. Cell elongation is not directly related to the radius of the body *a*, because of the different configuration of cells between asexual colonies and sperm packets (Figure 9). Elongated cells in sperm packets are hemispherically aligned with all flagella anterior, whereas spherical cells in asexual colonies are spherically aligned with all flagella out and with a hollow center.

**Table 1.**
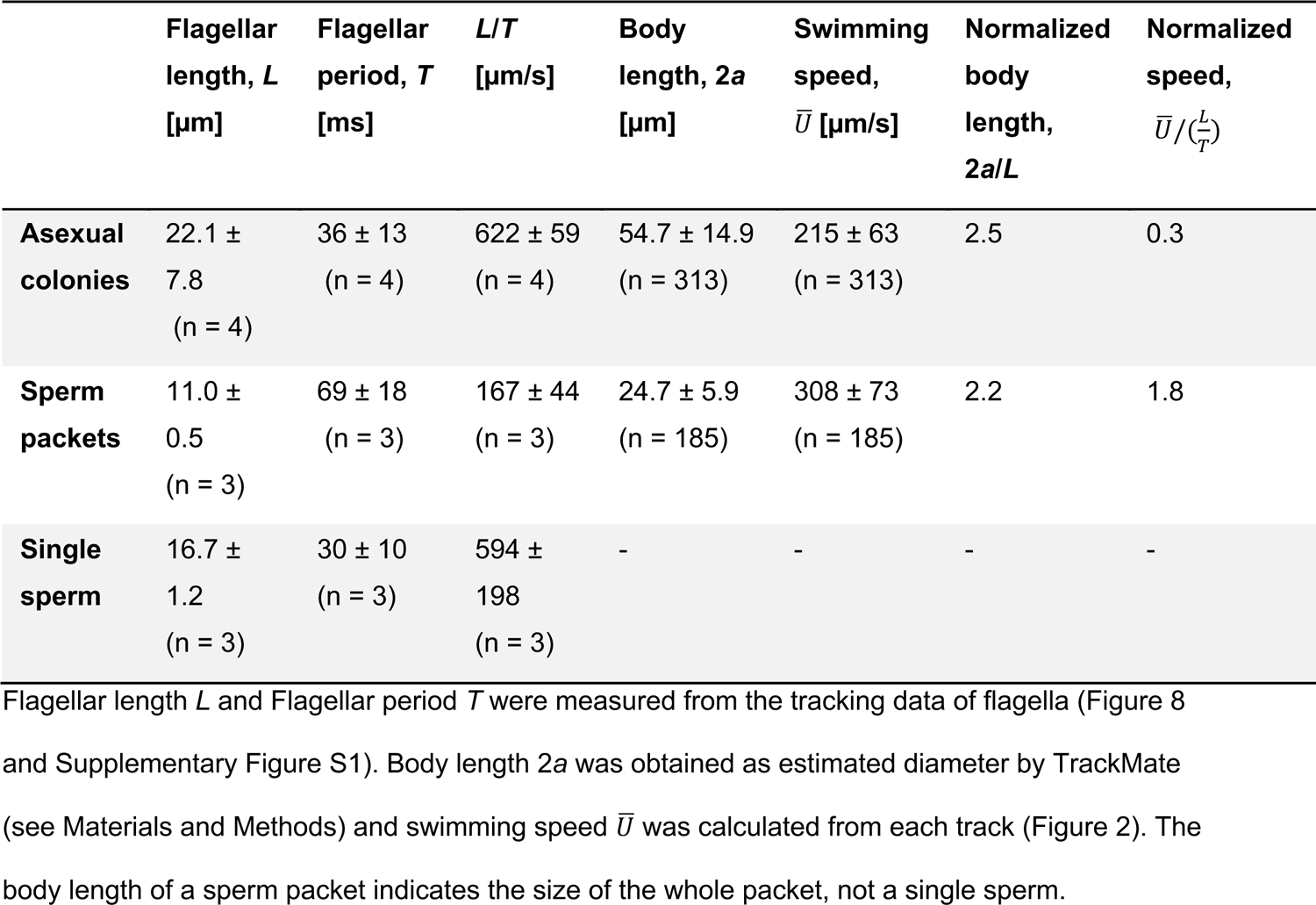
Swimming parameters of asexual colonies, sperm packets, and dissociated single sperm.

Although the beat period of sperm packets was longer than that of asexual colonies (i.e., the flagellar beat was slower in sperm packets), swimming was faster in sperm packets than in asexual colonies. This could be attributed to the differences in size and distribution of flagella between sperm packets and asexual colonies as well as the difference in the waveform and collective motion of the flagella. The effect of size difference (i.e., sperm packets were smaller in size and thus had smaller drag than asexual colonies) might be stronger than the effect of the beat period difference. Another reason might be the differences in flagellar motion. Although flagellar movements of both sperm packets and asexual colonies were categorized as asymmetric motion, they showed subtle differences in actual waveforms (Figure 8A and C) and spatiotemporal dynamics in curvatures (Figure 8D and F). These differences, as well as the differences in metachronal wave dynamics (Figure 4B, Figure 7B) and hydrodynamic interactions between flagella, would result in faster swimming in sperm packets.

### Switching mechanism of symmetric and asymmetric waveforms

We found that the flagellar waveform changed in sperm packets and dissociated single sperm. Sperm packets showed an asymmetric waveform, whereas dissociated single sperm showed a symmetric waveform. Wild-type *Chlamydomonas* flagella usually show an asymmetric waveform, and sometimes the waveform becomes symmetric, depending on the Ca^2+^ concentration inside the flagella [44,45]. A similar mechanism can be considered for waveform switching in the sperm of *P. starrii*: the Ca^2+^ concentration is constantly higher in the flagella of dissociated single sperm than in the sperm packets. To test this hypothesis, an experiment using demembranated models of both sperm packets and dissociated single sperm is desirable. In animal sperm, Ca^2+^ plays an important role in changing flagellar waveform (and, thus, trajectory) in response to chemoattractants [46]. It is possible that Ca^2+^ may also play a role in the reproduction of volvocine green algae.

### Flagellar lengths in dissociated single sperm and sperm packets

From the tracking data of flagella (Figure 8), we found differences in the lengths of flagella for sperm packets and dissociated single sperm. The length was 16.7 ± 1.2 µm in single dissociated sperm (n = 3) and 11.0 ± 0.5 µm in sperm packets (n = 3). As the sample size was too small to perform statistical tests, we only discuss these data from the viewpoint of descriptive statistics. The mean length was approximately 6 µm longer in single dissociated sperm than in sperm packets.

Although the difference might be partly attributed to the three-dimensionality of flagellar waveform especially in sperm packets, the flagella of the fixed sperm packet also looked short (Figure 9A). The flagella might be elongated before dissociation of sperm packets; that is, sperm might undergo a process of maturation during which flagella might grow. In *Chlamydomonas*, the growth rate of flagella after deflagellation is reported to be approximately 9 µm/hour [47]. Thus, it would be possible for the *Pleodorina* sperm flagella to grow approximately 6 µm on a timescale of 1 hour before dissociation. Currently, it is difficult to control the process of dissociation (i.e., the age of sperm packets), but future study should test the maturation hypothesis of sperm before sperm packet dissociation.

### Significance of faster swimming speed in sperm packets

The sperm packets swam faster than the asexual colonies. To date, we have not measured the swimming speed of female sexual colonies, which also have flagella. However, it would be reasonable to assume that the speed of female sexual colonies would not be significantly higher than that of asexual colonies because the morphology does not differ significantly between sexual and asexual colonies of females [24]. A previous study reported that female colonies of *Eudorina elegans*, a close relative to *Pleodorina starrii*, were immobile [48]. Taking the safe side, if we consider that the speed of female sexual colonies is not significantly different from that of asexual colonies of male strains, the faster swimming of sperm packets would enable them to catch up with female colonies during sexual reproduction. However, it remains unclear how the sperm packets approach female colonies. Three possibilities can be considered: accidental proximity, hydrodynamic attraction as discussed in Drescher et al. [49], and chemical attraction as occurs in some other taxa. Although chemoattractants that induce alterations in sperm flagellar waveforms have been identified in some species of animals [46] and brown algae [50], such an attractant is unknown in volvocines. Examining chemotaxis and determining chemoattratants, if any, in *P. starrii* will be the focus of a future study.

## Conclusions

In this study, we quantitatively characterized the swimming and flagellar motility of three forms of the volvocine green alga *Pleodorina starrii*, with emphasis on comparison between asexual colonies and sperm packets. Two important changes in the flagellar properties during male gametogenesis and gamete fusion were confirmed. The first is the change in flagellar distribution during sperm packet formation, which allows the packets to approach the female sexual colony at high speed. This change in flagellar distribution is accompanied by a drastic change in swimming mode, from neutral swimmers (asexual colonies) to pullers (sperm packets). The second is the change from asymmetric to symmetric waveform for effective fusion with the female gametes. These findings revealed diverse, developmentally- and physiologically-controlled movements under low Reynolds number conditions even in a male strain of one species. Further clarification of such diverse movements will lead to the control of motion and cargo transport under low Reynolds number conditions. Further biophysical, biochemical and bioinformatics studies should clarify the phototaxis and chemotaxis of sperm packets, the mechanism of switching flagellar waveforms, and the entire process of sexual reproduction in the context of swimming.

## Acknowledgements

This study was supported by JSPS KAKENHI #21K20661 (A.K.) and #22H05689 (A.K.). We thank Dr. Masafumi Hirono (Hosei University) for the immunofluorescence staining protocol, Ms. Tomoka Sakamoto (Narishige) for technical help with the suction experiments, Dr. Toshihiro Omori (Tohoku University) for discussions on fluid mechanics, Dr. Masato S. Abe (Doshisha University) for discussions on statistical methods, and Dr. Muneyoshi Ichikawa (Fudan University) for advice on revision of figures.

## Author contributions

A.K. conceived the study. K.T., H.N., and T.H. prepared the samples. T.N. constructed the microscopy system. S.A.B. provided the software and developed the image analysis methods. A.K. performed the experiments. A.K. and S.A.B. analyzed the data. A.K. drafted the manuscript, and A.K., K.T., H.N., and T.N. contributed to the final manuscript.

## Supporting Information Files

**S1 File.** Supplementary methods, figures, and tables.

**S1 Movie.** Supplementary Movie S1. Sperm packets viewed from the side using dark-field microscopy (objective ×10). The sperm packets swam along the anterior flagella. Playback speed, 1/50.

**S2 Movie.** Supplementary Movie S2. Movement of flagella in an asexual colony in dark-field microscopy (objective ×10). A movie of a rolling colony near the wall was recorded, as described in the Supplementary Methods, and the registered colony is shown. Playback speed, 1/50.

**S3 Movie.** Supplementary Movie S3. Dissociation of sperm packets observed using dark-field microscopy (objective ×10). Playback speed, ×2.

**S4 Movie.** Supplementary Movie S4. Single dissociated sperm observed using dark-field microscopy (objective ×10). Playback speed, 1/50.

**S5 Movie.** Supplementary Movie S5. Sperm packets were held with a micropipette and observed using phase-contrast microscopy (objective ×40). Contrast enhanced. Playback speed, 1/50.

## Notes

https://zenodo.org/records/10724959

### Competing Interest Statement

The authors have declared no competing interest.

### Summary of Updates

Texts and figures were thoroughly revised. Part of flagellar tracking in Figure 8 and S1 was improved. The scale bar in Figure 9A was corrected. Two subsections are added to the Discussion section.

