## Supplementary Methods, Figures and Tables for "Swimming ability and flagellar motility of sperm packets of the volvocine green alga *Pleodorina starrii*"

### Analysis of metachronal waves in asexual colonies

To analyze metachronal waves in asexual colonies (Figure 4), movies of rolling colonies observed using dark-field microscopy were registered (i.e., rotation and translation were corrected, and the colony in the image was maintained at a constant position and angle) using the “Register Virtual Stack Files” Plugin of Fiji with the “Rigid” setting. For proper registration of the colony, images of the wall near the colony were deleted and filled with black using the “Edit > Fill” command of Fiji. Kymographs of the movement of flagella were drawn by segmented lines using the “Image > Stacks > Reslice” command.

### Supplementary Figures

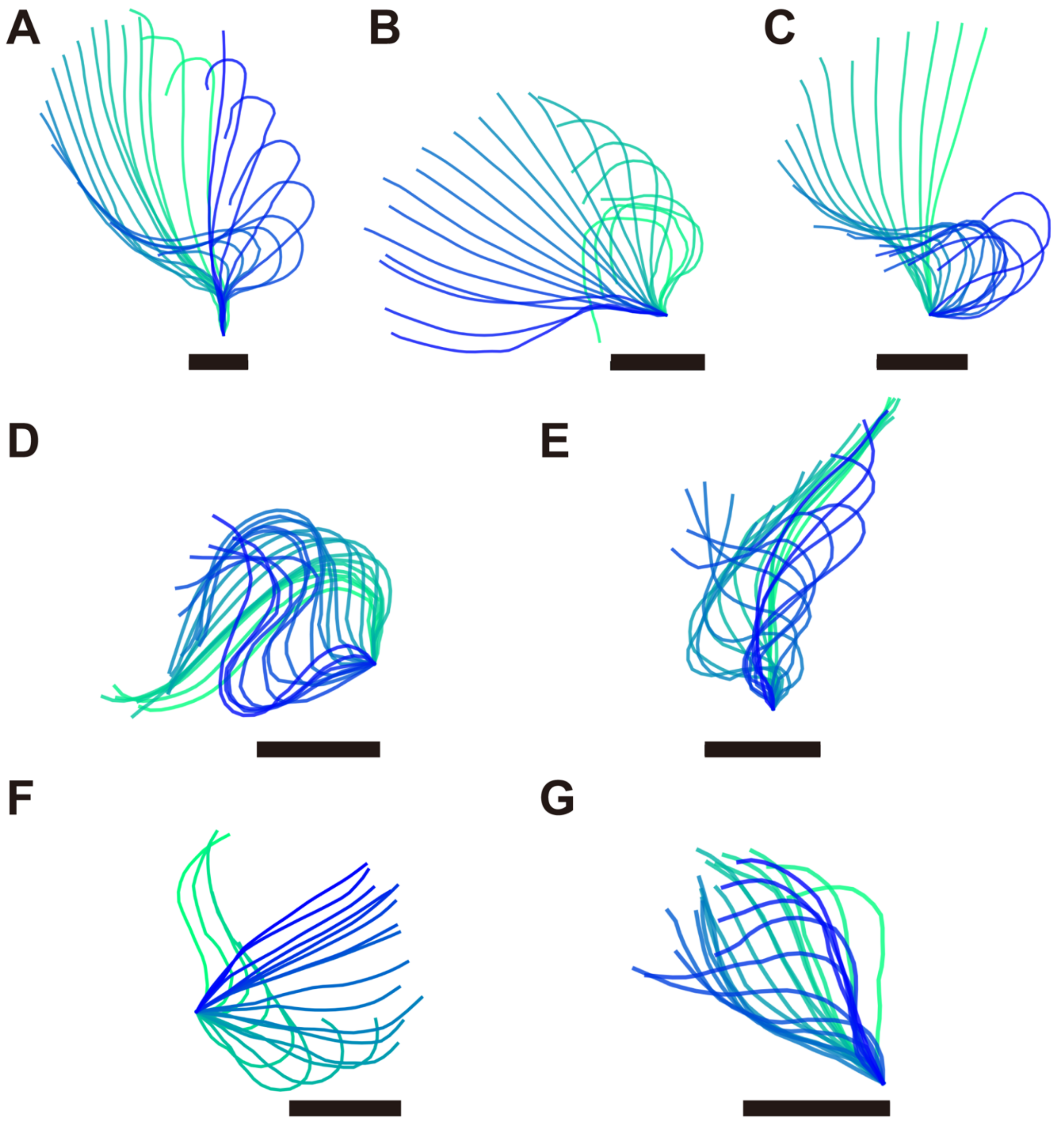


Supplementary Figure S1. Traces of flagella. (A)–(C) Asexual colonies, (D)–(E) single dissociated sperm, and (F)–(G) sperm packets. Scale bars, 5 μm. Original movies were taken at 1,000 fps. In (A), (E), and (F), 60 frames are shown every 3 frames. Twenty frames are presented in (B)–(D). In (G), 100 frames are shown every five frames. The flagellar lengths listed in Table 1 and Supplementary Table S1 were measured from these data and the data shown in Figure 8.

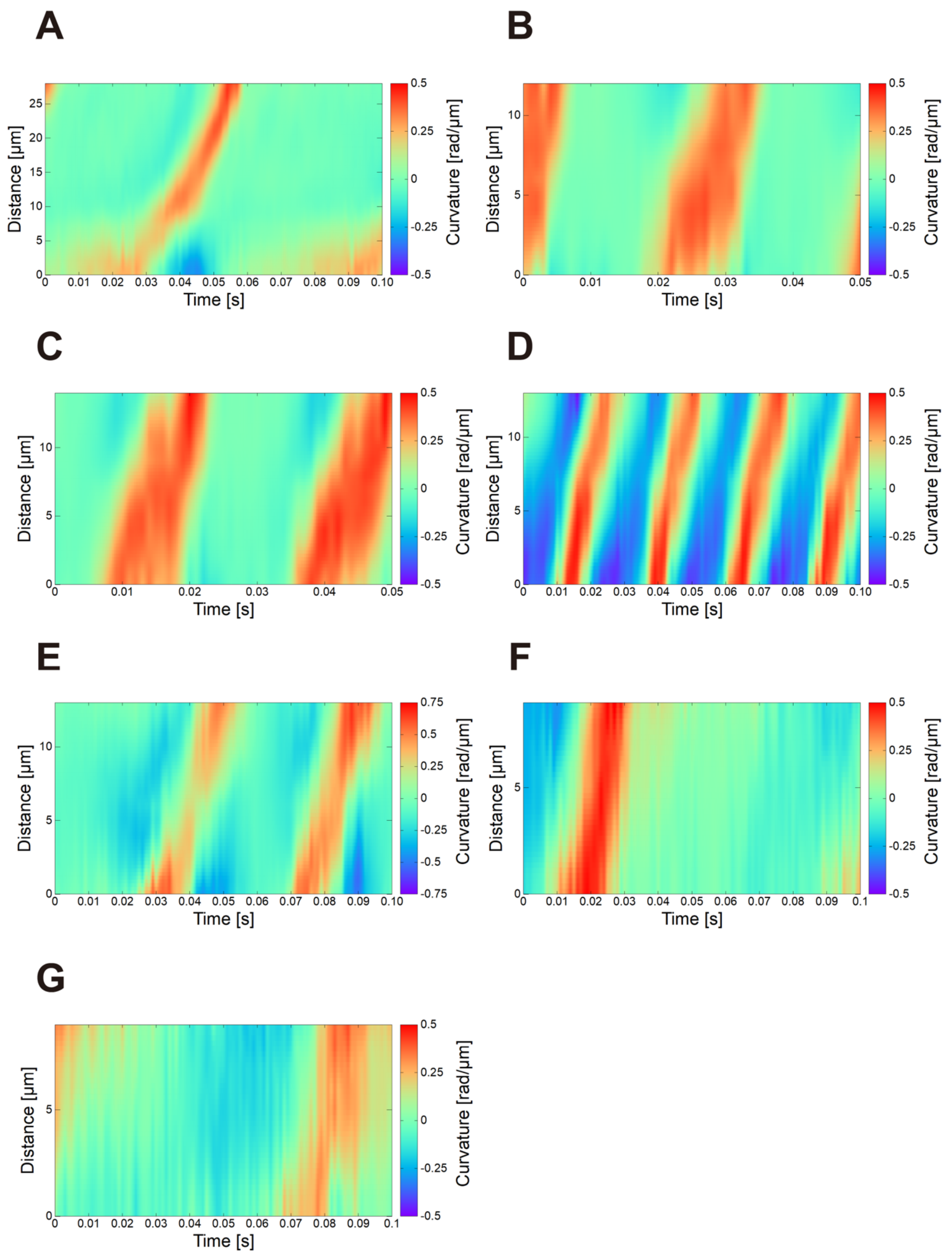


Supplementary Figure S2. Space-time plots of curvature along with flagellar length and time. (A)–(C) Asexual colonies, (D)–(E) single dissociated sperm, and (F)–(G) sperm packets. The beat periods listed in Table 1 and Supplementary Table S1 were estimated from these data and the data shown in Figure 8.

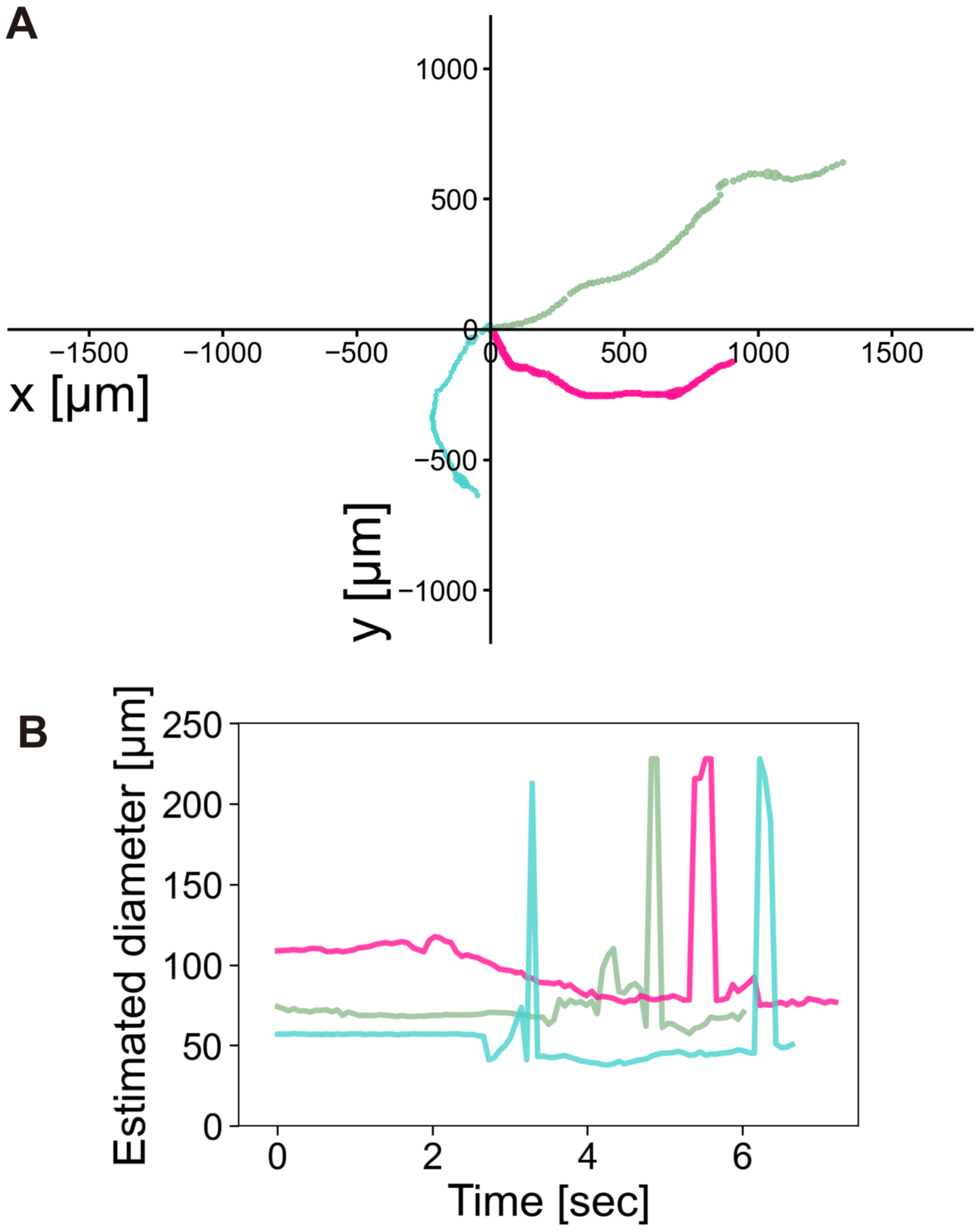


Supplementary Figure S3. Examples of estimated diameter along the trajectories. (A) Three trajectories of asexual colonies. The size of points represents estimated diameter. (B) Time course of estimated diameter. Peaks were observed when the objects were out of focus. Colors correspond to trajectories in (A).

### Supplementary Tables

Supplementary Table S1. Raw flagellar motility parameters. All data for beat period and flagellar length are shown. Flagellar length shows the mean ± standard deviation from 50–100 frames.

| **ID** | **Category** | **Flagellar length, L [μm]** | **Beat period, T [ms]** | **L/T [μm/s]** | **Trace** | **3D plot** |
|---|---|---|---|---|---|---|
| **1** | Asexual colony | 33.43 ± 1.57 | 55.5 | 602.0 | Fig. S1A | Fig. S2A |
| **2** | Asexual colony | 21.23 ± 2.11 | 29.9 | 709.8 | Fig. 8A | Fig. 8D |
| **3** | Asexual colony | 17.04 ± 1.31 | 28.9 | 590.2 | Fig. S1B | Fig. S2B |
| **4** | Asexual colony | 16.65 ± 1.01 | 28.4 | 586.2 | Fig. S1C | Fig. S2C |
| **5** | Single sperm | 17.94 ± 0.94 | 24.2 | 740.2 | Fig. S1D | Fig. S2D |
| **6** | Single sperm | 16.62 ± 0.95 | 24.7 | 673.0 | Fig. 8B | Fig. 8E |
| **7** | Single sperm | 15.60 ± 1.07 | 42.3 | 368.6 | Fig. S1E | Fig. S2E |
| **8** | Sperm packet | 11.49 ± 0.65 | 80.8 | 142.3 | Fig. S1F | Fig. S2F |
| **9** | Sperm packet | 11.10 ± 0.65 | 78.3 | 141.7 | Fig. S1G | Fig. S2G |
| **10** | Sperm packet | 10.51 ± 1.33 | 48.3 | 217.8 | Fig. 8C | Fig. 8F |

### Supplementary Movies

Supplementary Movie S1. Sperm packets viewed from the side using dark-field microscopy (objective ×10). The sperm packets swam along the anterior flagella. Playback speed, 1/50.

Supplementary Movie S2. Movement of flagella in an asexual colony in dark-field microscopy (objective ×10). A movie of a rolling colony near the wall was recorded, as described in the Supplementary Methods, and the registered colony is shown. Playback speed, 1/50.

Supplementary Movie S3. Dissociation of sperm packets observed using dark-field microscopy (objective ×10). Playback speed, ×2.

Supplementary Movie S4. Single dissociated sperm observed using dark-field microscopy (objective ×10). Playback speed, 1/50.

Supplementary Movie S5. Sperm packets were held with a micropipette and observed using phase-contrast microscopy (objective ×40). Contrast enhanced. Playback speed, 1/50.
